# Increasing temperature threatens an already endangered coastal dune plant

**DOI:** 10.1101/2020.08.02.233288

**Authors:** Aldo Compagnoni, Eleanor Pardini, Tiffany M. Knight

## Abstract

Climate change has the potential to reduce the abundance and distribution of species and threaten global biodiversity, but it is typically not listed as a threat in classifying species conservation status. This likely occurs because demonstrating climate change as a threat requires data-intensive demographic information. Moreover, the threat from climate change is often studied in specific biomes, such as polar or arid ones. Other biomes, such as coastal ones, have received little attention, despite being currently exposed to substantial climate change effects. We forecast the effect of climate change on the demography and population size of a federally endangered coastal dune plant (*Lupinus tidestromii*). We use data from a 14-year demographic study across seven extant populations of this endangered plant. Using model selection, we found that survival and fertility measures responded negatively to temperature anomalies. We then produced forecasts based on stochastic individual based population models that account for uncertainty in demographic outcomes. Despite large uncertainties, we predict that all populations will decline if temperatures increase by 1° Celsius. Considering the total number of individuals across all seven populations, the most likely outcome is a population decline of 90%. Moreover, we predict extinction is certain for one of our seven populations. These results demonstrate that climate change will profoundly decrease the current and future population growth rates of this plant, and its chance of persistence. Thus, our study provides the first evidence that climate change is an extinction threat for a plant species classified as endangered under the USA Endangered Species Act.

## INTRODUCTION

Year to year variation in weather influences demographic processes such as survival, growth and reproduction, and ultimately population persistence (Sæther et al. 2000). For many natural populations, climate change will increase the frequency of weather conditions outside the physiological tolerances of individuals, and is thus expected to reduce the number, size and distribution of populations (IPCC 2014). As a result, conservation planning (Akçakaya et al., 2014) and projections of biodiversity change (Pereira et al. 2010) will benefit from understanding how climate change will threaten population viability.

Researchers typically use species distribution models (SDMs) to assess the effects of climate change on species (Elith and Leathwick 2009). However, species distributions might include declining populations of the species because of a currently unsuitable climate (Schurr et al. 2012). Subsequently, while SDMs describe current ranges well, their prediction of future ranges are highly uncertain (Keenan et al. 2011). Hence, forecasting the effect of climate change on species persistence should be more reliable using process-based models that require long-term demographic data across the entire life cycle of the organism (Ehrlén and Morris 2015, Paniw et al. 2019). Despite this, our knowledge on how climate drives demography is relatively limited, with most studies coming from animals in cold and arid biomes. For example, changing temperature has been linked to reductions in population growth rates of alpine plants (Doak and Morris 2010, Campbell 2019, Iler et al. 2019) and polar animals (Jenouvrier et al. 2009, Hunter et al. 2010). Changing precipitation has been linked to changes in population growth rates of organisms in arid environments (Tews and Jeltsch 2004, Jonzén et al. 2010). However, the effect of climate change in other biomes, regardless of their extent or importance to humans, has received less attention.

Coastal ecosystems are a biome for which the extinction threat from climate change has received little attention. This occurs despite the importance of species conservation for human well-being (Díaz et al. 2019), and despite the importance of coastal ecosystems and their susceptibility to climate change. Coastal ecosystems occupy a small portion of terrestrial surface, but are home to 41% of the human population (Martínez et al. 2007) and to a large portion of the earth’s economy; for example, 31% of the USA gross domestic product in 1985 (Luger 1991). Coastal biomes protect inland ecosystems during storms by attenuating or resisting wave action (Barbier 2015). Current and future climate change in coastal ecosystems is altering precipitation patterns, increasing the frequency of high temperature extremes (USGCRP 2018), and causing habitat loss and salt water inundation (Feagin et al. 2005). To date, demographic research on coastal dune plants has focused on understanding the effects of plant-animal interactions and succession (e.g. Maron 1998, Dangremond et al. 2010a, Pisanu et al. 2012, Pardini et al. 2015, Cogoni et al. 2019). Thus, it is timely to consider the effects of climate on the demography of coastal species.

Understanding the effect of climatic drivers on future population size requires quantifying how the link between climate and demography translates into population dynamics. Studies establishing such link are the foundation to prioritize conservation efforts. For example, the polar bear was listed as a threatened species under the endangered species act thanks to demographic studies (USFWS 2015). Demographic analysis linked warming-induced reductions in sea ice extent to reductions in survival and breeding probabilities, and increased risk of extinction (Hunter et al. 2010). As a result of this work, the US Fish and Wildlife Service indicated curbing Arctic warming as the most important conservation action for this species (USFWS 2015). Such demographic studies are most useful when carried out across multiple populations, in order to account for the substantial variation in populations sizes and vital rates (Glenn et al. 2010).

Here, we examine the effect of changing climate on the vital rates and population growth of a perennial coastal dune plant. We first link the effect of climate on separate vital rates, and we use an integral projection model to estimate their contribution to population growth rate. Then, we use an individual based model to forecast the effect of projected climate change on future population numbers in the next 30 years. These numerical simulations account for the effect of initial population numbers, year to year stochastic weather, and demographic stochasticity. We parameterize these models using a 14-year data set that comprehensively sampled 7 of the 15 known remaining populations of this species. The populations we included range from the smallest to the largest known (17 and 180000 individuals, respectively). Our focal species, *Lupinus tidestromii*, is listed as endangered under the USA endangered species act, but to date, climate change is not considered a threat to this or any plant species.

## MATERIALS AND METHODS

### Study Species and Site

*Lupinus tidestromii* (Fabaceae) is a perennial, herbaceous plant that is endemic to coastal dunes in northern California, where it is found in 15 extant populations. These populations are located in Sonoma, Monterey, and Marin counties, seven of these are contained within Point Reyes National Seashore (Figure 1). In this protected area, the population located at Abbotts Lagoon contains more than 50% of the remaining individuals of the entire species (USFWS 2009). *L. tidestromii* was listed as federally endangered in 1992 and among its primary threats are habitat loss, trampling by visitors and cattle, hybridization, direct competition with invasive plants, and elevated levels of seed predation in the presence of the invasive grass *Ammophila arenaria* (USFWS 2009, Dangremond et al. 2010b). Our study was conducted at Point Reyes National Seashore (latitude: 33.1120, longitude: −122.9579). This site has a Mediterranean climate, with wet winters and dry summers (Evens 2008).

**Figure 1.**
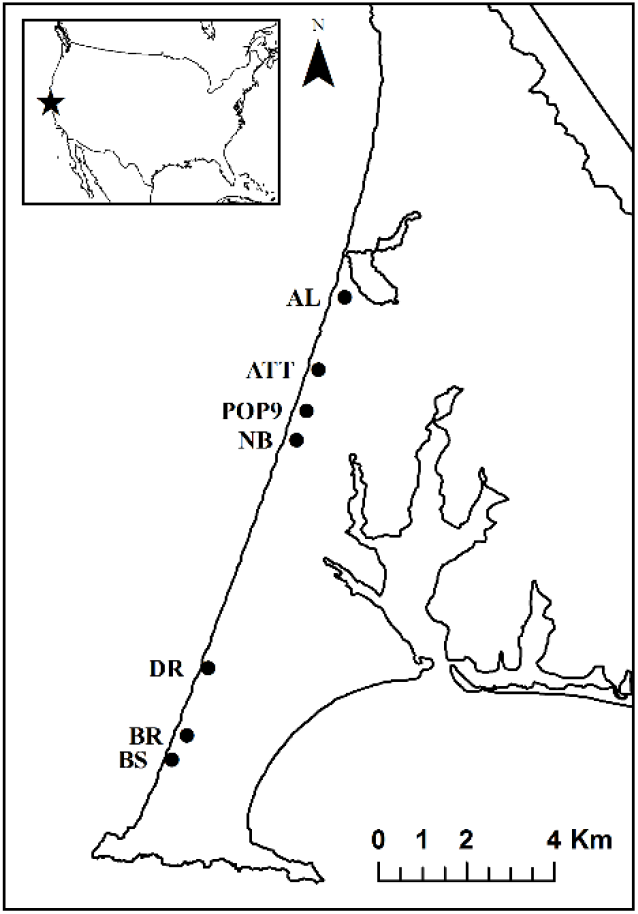
Map of the seven study populations of *L. tidestromii* at Point Reyes National Seashore, Marin County, California, USA. The entire range of this species includes two additional clusters of populations located in Sonoma and Monterey county.

*L. tidestromii* is a herbaceous perennial plant that typically lives three years. It produces prostrate stems, and small, upright inflorescences with a whorl of flowers that produce leguminous fruits (Baldwin et al. 2012). Seeds are dispersed locally by explosive dehiscence (USFWS 1998). Seeds have a tough coat that likely allows survival for many years in the seed bank. However, seeds that do not germinate within the first two years become buried by sand. We assume these seeds do not contribute to the short-term population dynamics, but that they can contribute to population growth after disturbance (Pardini et al. 2015). Flowers experience occasional insect herbivory, and fruits and seeds experience pre- and post-dispersal seed predation by the native *Peromyscus maniculatus* (deer mouse). We have never observed pre-dispersal insect seed predation.

### Monitoring

We began demographic censuses in 2005 at three sites (AL, ATT, NB) in 2005, and expanded monitoring at an additional four sites (Pop9, DR, BR, and BS) in 2008 (Fig. 1). Populations NB, BS, and DR are small enough that we monitored every individual plant in the population each year. In AL and BR, and ATT and Pop9 starting in 2013 and 2015, we stratified sampling, censusing plants located in clusters across the extent of the population. We haphazardly chose plants across a range of sizes and microhabitats. Annually, we also tagged at least 50 new seedlings across each site. We know total population counts for ATT, Pop9, NB, BS, and DR for every year. We inferred population counts for NB, BS, and DR, and we counted the total number of individuals in the populations after starting stratified sampling in ATT and Pop9.

### Vital rate data

To quantify the vital rates, in 2005, we started censusing tagged plants in June when fruits on most plants were dehiscing. These censuses quantified the transition rates from one year (*t*) to the next (*t+1*). To track individuals, we attached numbered aluminum tags to the basal stem of plants, or staked these tags near plants. We relocated plants each year using a GPS and a metal detector. For each plant, every year we recorded size, survival, flowering status, and number of flowering inflorescences. We quantified size approximating the surface area covered by each individual (Appendix S1).

For reproductive plants, we recorded the total number of inflorescences and of those that were aborted, consumed, or intact. Flowers on inflorescences can abort or produce fruits; fruits can be clipped by mice or remain intact, dehisce and disperse seeds. We used the fate of inflorescences to calculate abortion, and consumption (pre-dispersal seed predation), for each population and year.

To calculate abortion, we summed the number of aborted inflorescences and divided by the sum of the total number of observed inflorescences. To calculate consumption, we summed the number of clipped inflorescences, and divided by the sum of the number of not aborted inflorescences. The resulting abortion and consumption are population-level means, because we performed the sums to calculate these metrics across all the plants in each population. We do not have observations on abortion and consumption for all populations and all years. We have abortion rates for all populations in 2010 and 2011, and from 2013 to 2017 (Appendix S2: Fig. S2) and consumption rates for all years except 2012 (Appendix S2: Fig. S3). In years and locations for which we did not have an abortion or consumption rate, we used the population-specific mean taken across censuses.

We assumed the number of fruits per inflorescence, and the number of seeds per fruit are constant across years and populations. We calculated fruits per inflorescence using data from 2011, averaging the number of fruits across all inflorescences for each plant, and then averaging across plants to produce a population-level value. We quantified the average number of seeds per fruit by counting seeds in 213 randomly collected fruits from multiple years and populations: AL: 2005 (23), 2008 (3), 2009 (15), 2010 (39); ATT: 2011 (33); BR: 2010 (23); DR: 2010 (10), 2011 (17); NB: 2010 (16) Pop9: 2010 (15).

We estimated germination rates through a field germination trial and calculated a recruitment adjustment factor to account for post-dispersal seed predation, and for other sources of unaccounted seed or seedling mortality. We installed caged seed baskets in 2008 at the AL population, and recording the proportion of seedlings that germinated in 2009 (*g*_1_), 2010 (*g*_2_), and 2011 (*g*_3_) (Pardini et al. 2015). Here *g*_2_, and *g*_3_ represent emergence out of seed bank stages. We then calculated a recruitment adjustment factor to account for post-dispersal seed predation and for other sources of unaccounted seed or seedling mortality. To account for these other sources of seed and seedling loss, we used our known population counts to estimate a recruitment adjustment factor, δ_*p*_, for each *p* population (Appendix S3). We modified germination rates *g*_1_, *g*_2_, and *g*_3_ employed in the IPM using δ_*p*_ as *g*_*t*_ (1 − δ_*p*_). We could only calculate δ_*p*_ for the five populations with known population counts (see “monitoring”). For population BR and ATT, we applied the mean of *δp* from BS and DR, and from ATT and Pop9, respectively the closest populations to BR and ATT.

### Modeling of size-dependent vital rates and climate effects

We modeled four vital rates as size dependent processes for each year and site: survival, growth, the probability of flowering, and the number of inflorescences. We modeled these vital rates based on generalized mixed linear models. We fit models in R using the lme4 package (Bates et al. 2015), and we performed a preliminary model selection based on Akaike Information Criterion (Burnham and Anderson 2002). In this preliminary model selection, we compared models predicting vital rates using linear, quadratic, and cubic predictors. Our baseline model structure included a random intercept and slope (the effect of size on the vital rate) for both years and populations:

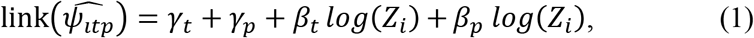

where 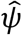 is the mean prediction of the vital rate model for individual *i*, in year *t*, at population *p*, *Z* is the size of individual *i*, the *γ* coefficients are normally distributed random intercepts, the β coefficients are normally distributed random slopes of log size, and *link* is the link function for the vital rate being modelled. The link function depended on the type of response variable. The models on survival (S) and flowering (F) used a logit link, because the response variable is Bernoulli distributed:

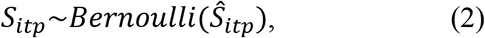

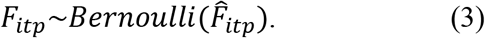

The number of inflorescences (R) used a log link, because this response variable follows a Poisson distribution:

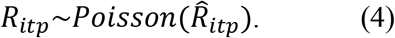

Finally, the growth model used an identity link: that is, it did not use a link function:

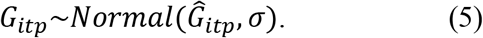

The best survival model was a cubic function of log(size), so that:

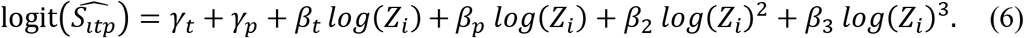

For the remaining three vital rates, the best model included only a linear predictor (Eq. 1). We selected which climatic predictor, if any, affected vital rates by performing a leave-one-year-out cross-validation. We selected whether climatic drivers increased the predictive power of our linear models (Eq. 1 and 6) and, if so, what climatic drivers improved predictive power the most. The models in this cross-validation simply added the climatic predictor to the vital rate models (Eq. 1–6). We considered four climatic predictors: temperature, precipitation, Oceanic Niño Index (ONI) (CPC n.d.), and the standardized precipitation evapotranspiration index (SPEI, Vicente-Serrano et al. 2009). We obtained temperature and precipitation data starting in 1990 from the PRISM dataset (Daly et al. 1994), the ONI data from the internet site of the Climate Prediction Center of the National Oceanic and Atmospheric Administration (CPC n.d.), and we derived the SPEI using our precipitation and temperature data using the R package SPEI (Beguería and Vicente-Serrano 2017). We calculated the yearly climate anomalies for each one of these four drivers. We computed anomalies using the mean and standard deviation observed between 1990 and 2018. Here, we define “year” as the time elapsing between demographic censuses: from June to May of the following year.

For each of the four climatic drivers, we used two yearly anomalies: one referred to the year leading up to the current census (*t*), and one to the year leading up to the preceding census (*t-1*). For example, for year 2012 (t=2012) we considered the climate anomaly observed between June 2011 and May 2012, and between June 2010 and May 2011 to predict the flowering probability and number of inflorescences observed in June 2012, and to predict the survival and growth occurred from June 2012 to June 2013. We thus tested for a lag in the effect of climatic drivers on plant vital rates (see also Teller et al. 2016; Tenhumberg et al. 2018).

We measured model performance summing the negative log-likelihood of each out-of-sample prediction, with the best model having the lowest score. We used negative log-likelihood as it provides a scoring rule across all of our generalized linear models that yields maximum values when the predicted values coincide with the true ones (Gneiting and Raftery 2007). We fit the final models using the best climatic predictor (if any).

### Population projections

We quantified the effect of a changing climate on the population dynamics of *L. tidestromii* looking at its projected long-term population growth rates and population sizes. We projected long-term population growth rates using an integral projection model (IPM, Ellner et al. 2016), and future population sizes using an individual based model (IBM). We used IBMs to add demographic stochasticity, which is a key component to predict population sizes, particularly for smaller populations (Lande 1993, Caswell 2001).

We used an IPM because it is a computationally inexpensive way to perform stochastic population projections of species whose dynamics depend on a continuous state variable: in this case, the size of individuals. We constructed the IPM using the parameters of the vital rates models described above. We designed most of these IPM parameters to change based on year and population. The IPM projects the number of z-sized *L. tidestromii* individuals, represented by vector *n(z)*, to the number of z-sized individuals next year, represented by vector *n(z’)*. This population is also composed of seeds that germinate one year after the projection interval, represented by scalar *B1*, and two years after, represented by scalar *B2*. The dynamics of the population are given by

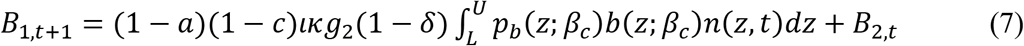

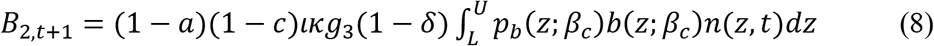

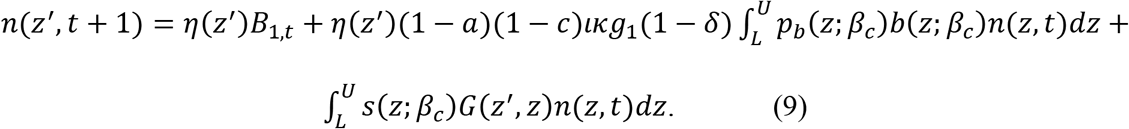

We show a schematic representation that links these equations to the life cycle of *L. tidestromii* in Figure 2. The equations above contain four vital rates that depend directly on the size of individuals, *z*, three of which depend on temperature anomaly, *β*_*c*_, while the remaining demographic rates are size- and climate-independent. Three vital rates are size- and climate-dependent: *p*_*b*_(*z*;*β*_*c*_) is the size- and climate-dependent probability of flowering, *b*(*z*;*β*_*c*_) is the size- and climate-dependent production of inflorescences, *s*(*z*;*β*_*c*_) is the size- and climate-dependent probability of survival, and *G(z’,z)* is size-dependent growth. The size- and climate-independent vital rates mostly refer to the transition from the number of inflorescences to the number of establishing seedlings. In particular, *a* is the population- and year-specific abortion rate, *c* is the population- and year-specific consumption rate (or pre-dispersal predation), *ι* is the average number of fruits produced per inflorescence, *κ* is the average number of seeds produced per fruit, δ is the recruitment adjustment factor (Appendix S3), and *g* values refer to germination rates. *g0* is the fraction of seeds that germinate before the end of the first transition, *g1* is the fraction of seeds that germinate before the end of the second transition, and *g2* is the fraction of seeds that germinate before the end of the third transition. Finally, *η(z’)* is the size distribution of seedlings emerging by the end of the transition.

**Figure 2.**
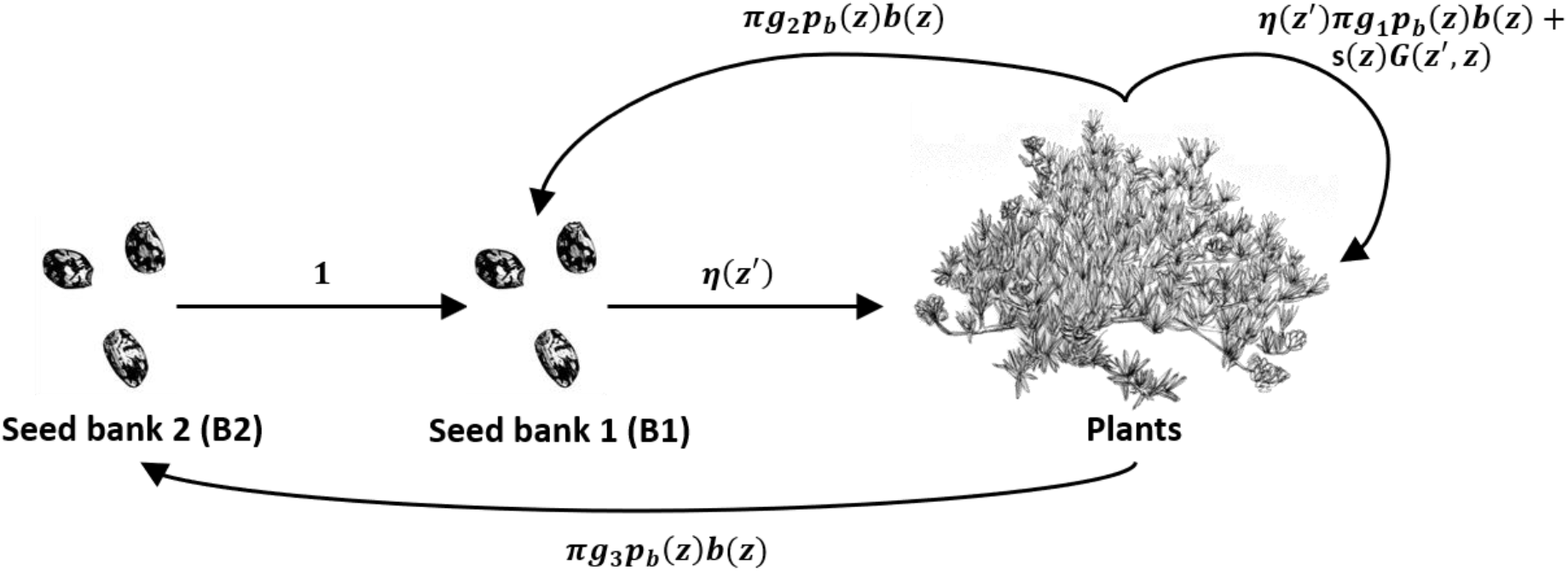
Life cycle model for *L. tidestromii*. This life cycle refers to the IPM described in Appendix S2 (Eq. 7–9). The IPM separates the population in three discrete stages: continuously sized plants (*plants*), seeds that germinate after one winter (*B1*), and seeds that germinate after two winters (*B2*). These stages are modeled, respectively, by equation 9, 7, and 8 in Appendix S2. To minimize the use of space, in this figure *π* corresponds to (1 − *a*)(1 − *c*)*lk*(1 − δ) in Equations 7–9.

We used this IPM to compute the effect of climate on λ_s_, the long-term stochastic population growth rate, and to elucidate this effect through perturbation analyses. We computed the stochastic population growth rate, λ_s_, by projecting the population numbers of each one of our seven population 50000 times. We started with a population of only one individual, discarding the first 1000 projections to allow populations to reach a stable size distribution. These stochastic simulations kept a constant average annual temperature, and randomly varied the year-specific conditions (parameters *γt* and *βt* in Eq. 1 and 6). We calculated λ_s_ as 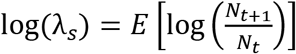, where E calculates the mean, and N is the total population size at the end (*t+1*) and at the beginning (*t*) of a transition. We repeated these projections across the whole range of observed climate anomalies we observed during our study period. Our best climatic predictor was always temperature. Therefore, we calculated λ_s_ from an average annual temperature of about 11° Celsius to an average annual temperature of 13°.

To understand the effects of temperature on λ_s_, we performed retrospective and prospective perturbation analyses (Caswell 2000). In particular, we performed a Life Table Response Experiment (LTRE, a retrospective analysis), and further analyzed its results through an elasticity analysis (a prospective analysis). The LTRE quantified the difference in λ_s_ observed when only one of the vital rates responded to deviations from the mean temperature. For example, we calculated λ_s_ for an IPM where only survival experienced the full range of temperature anomalies. In these simulations, all other vital rates experienced average climate (about 12° Celsius). We then calculated the contribution of each vital rate to the effect of climate. This contribution is the difference between the λ_s_ computed by these stochastic simulations, and the λ_s_ where all vital rates experienced a 12° Celsius climate.

To better interpret these LTRE results, we calculated vital rate elasticities for the lower level parameters of the IPM (Eq. 7–9; Fig. 2). The LTRE results depend on the sensitivity to the parameter **β*_*c*_* in the vital rate models (Eq. 7–9). However, we provided the elasticity analysis of all lower level parameters, to place these results in the context of the entire life cycle. We calculated these elasticities using the asymptotic population growth rate (λ) as a proxy for λ_s_.

### Forecast

We forecasted population numbers through an IBM that incorporates probabilistic events of the life cycle by projecting the dynamics of each individual. Given our life cycle (Fig. 2), these probabilistic events were: survival, growth, probability of flowering, production of inflorescences, the abortion of inflorescences, the consumption of inflorescences, and the germination of seeds. We simulated each of these processes using the appropriate statistical distributions (Appendix S4).

We used this IBM to produce a forecast of population sizes in the next 30 years that included uncertainty arising from process and demographic stochasticity. We projected each population for 30 time steps, and we replicated each run 1000 times. Each run included a different sequence of years, and a different realization of each vital rate process (Appendix S4, Eq. 1–7). We started each simulation with the number of individuals observed in June 2018, assuming the stable stage distribution suggested by the population-specific IPM model. We ran two simulations: one assuming the current mean climate (average annual temperature of 12° Celsius), and one assuming the largest weather anomaly we observed during our 2005-2018 study period (which was a year with an average temperature of 13° Celsius). A mean annual temperature of 13° Celsius is a conservative forecast, because projections suggest that in 2055, temperature will be on average 2.1 Celsius higher than the 1990-2018 mean (Vose et al. 2017). We used the 1000 simulations for each population and climate scenario to calculate 95% confidence intervals of population abundances during the 30-year projections.

## RESULTS

### Vital rates and population growth

Plant vital rates and consumption varied across our seven populations and 14-year study period (Appendix S2: Fig. S2–S7). As a result, asymptotic population growth rates (λ) was variable across space and time, and our two largest populations (AL, ATT) and the transitions years 2011-2012 and 2016-2017 were associated with relatively high values of λ (Appendix S2: Fig. S8). There was a remarkable population crash between 2014 and 2015 (Appendix S2: Fig. S8).

### Observed and future climate variation

Across our 14-year study period, we observe variation in weather on par with that across longer time periods (Appendix S5: Fig. S9), but not within the range of weather projected with climate change by the middle of the century (Wang et al. 2016, n.d.). For example, mean annual temperature was 11.5° C across the study period, 12° C from 1990-2018. This temperature is projected to increase to about 14° C (range: 13.8-14.6° C) by 2055 according to the Representative Concentration Pathway of 4.5 W/m^2^ (Wang et al. 2016, n.d., Vose et al. 2017). The range of annual temperatures observed from 1990-2018 was 11-13° C, and thus we only consider conservative scenarios of a 1° C increase in temperature.

### Vital rates and population responses to climate

Growth did not respond to climate, but the remaining three size-dependent vital rates correlated negatively with average annual temperature (Fig. 3, Appendix S2: Fig. S4, Fig. S6–7). There is a sharp decline in population growth rate with increasing mean annual temperature (Fig. 4A).

**Figure 3.**
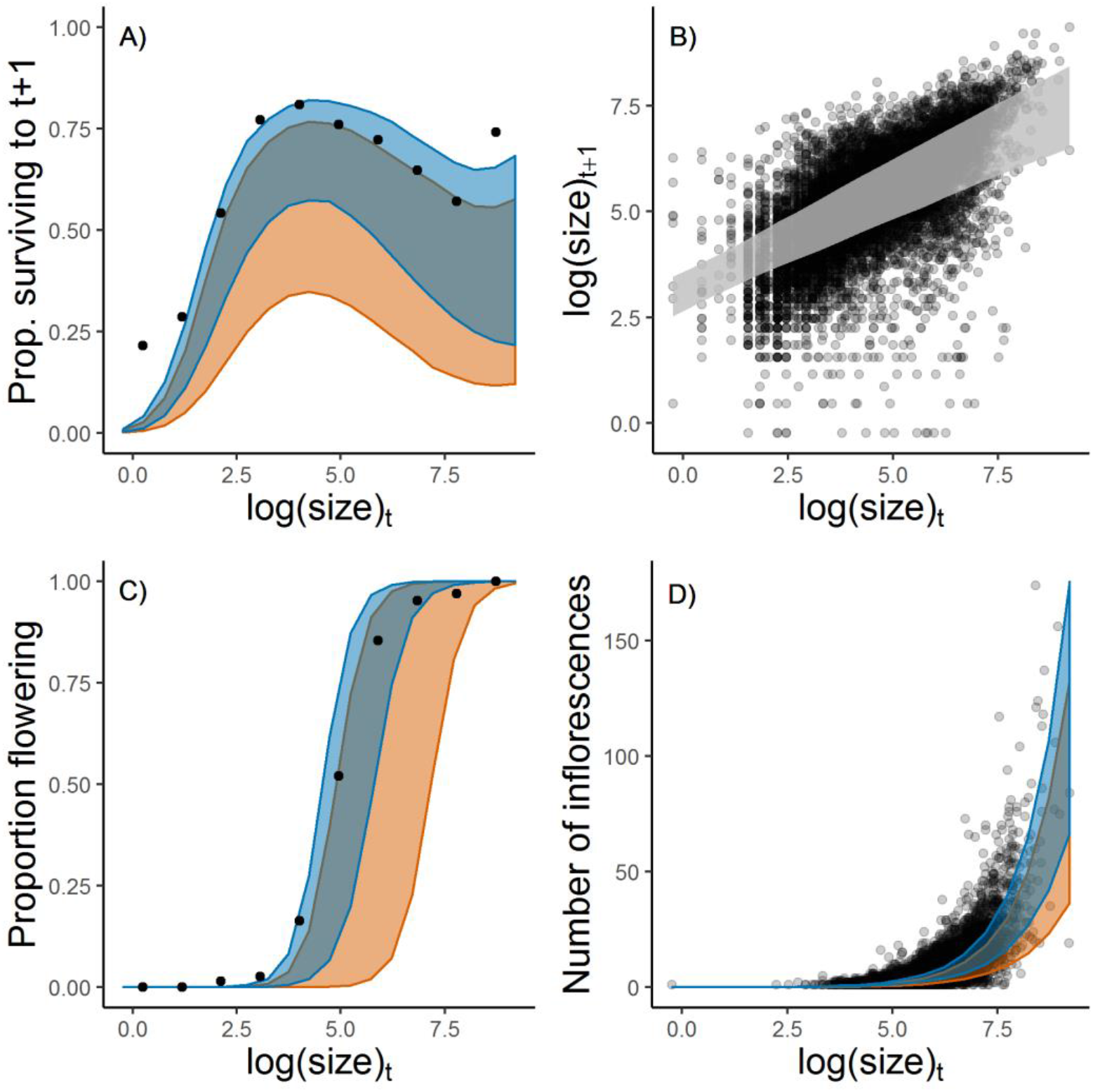
The effect of plant size and mean annual temperature (when relevant) on survival (A), growth (B), the probability of flowering (C), and the production of racemes (D). The dots in the plots showing survival (A) and probability of flowering (C) represent the average proportion of individuals surviving and flowering, respectively, within 10 equally spaced intervals of individual log sizes. Shaded areas show the 95% confidence intervals of the mean model predictions. For the three climate-dependent vital rates, we show credible intervals of vital rates responding to the average yearly temperature observed between 1990 and 2018 (12° Celsius, in blue), and the highest average yearly temperature observed during the same period (13° Celsius, in orange).

**Figure 4.**
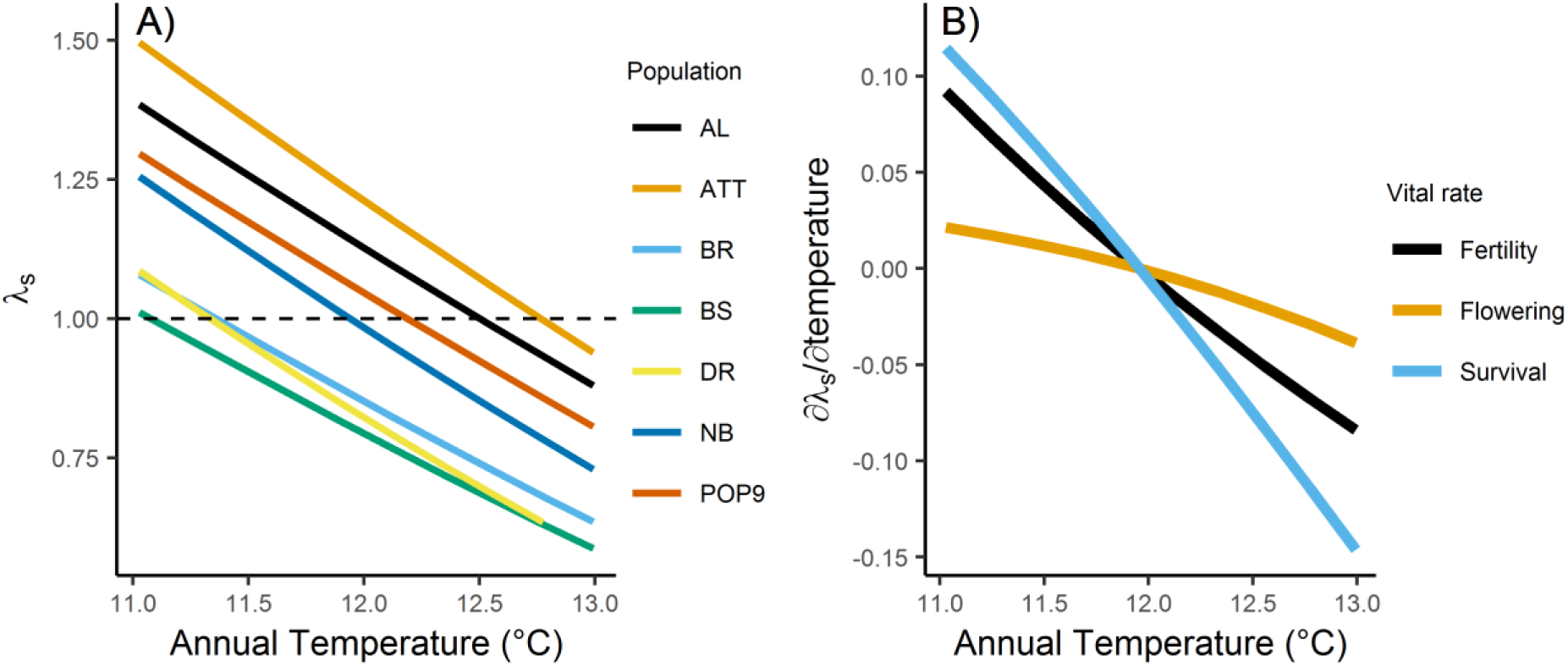
Long-term stochastic population growth rate (λ_s_) as a function of mean annual temperature for each of the seven *L. tidestromii* populations (A), and the partial effect of each climate-dependent vital rate on the λ_s_ (B). The average yearly temperature observed between 1990 and 2018 was 12° Celsius; the average temperature during the study period was 11.5° Celsius.

While the slope of the relationship between population growth rate and annual temperature is similar for all populations, these vary in their mean vital rates (i.e., their intercepts and slopes, Appendix S2: Fig. S4–7). If the temperatures remained at the 1990-2018 mean (12° C), three of the four smallest populations would be expected to decline. At these temperatures, the remaining four populations, including the three largest, are projected to grow. On the other hand, in a scenario where mean annual temperature increases by 1° C, all populations are projected to decline (Fig. 4A).

The LTRE analyses indicate that the decline in population growth rate with increasing temperature is primarily due to the effects of temperature on plant survival (Fig. 4B). Two factors explain the high contribution of plant survival on the change in population growth rate. First, population growth rate is very sensitive to changes in survival (Appendix S6: Fig. S10), and second, survival responds dramatically to increasing temperature (Fig. 4A).

### Forecast

Population trajectories indicate that if temperature remained at its 1990-2018 average, this endangered plant species would double its population number in the next 30 years. However, with a 1° C increase in temperature, there would be a 90% average reduction in individuals (Fig. 5). This scenario is conservative, as more dramatic increases in temperature are projected for this region in the next 30 years (Wang et al. 2016, n.d., Vose et al. 2017).

**Figure 5.**
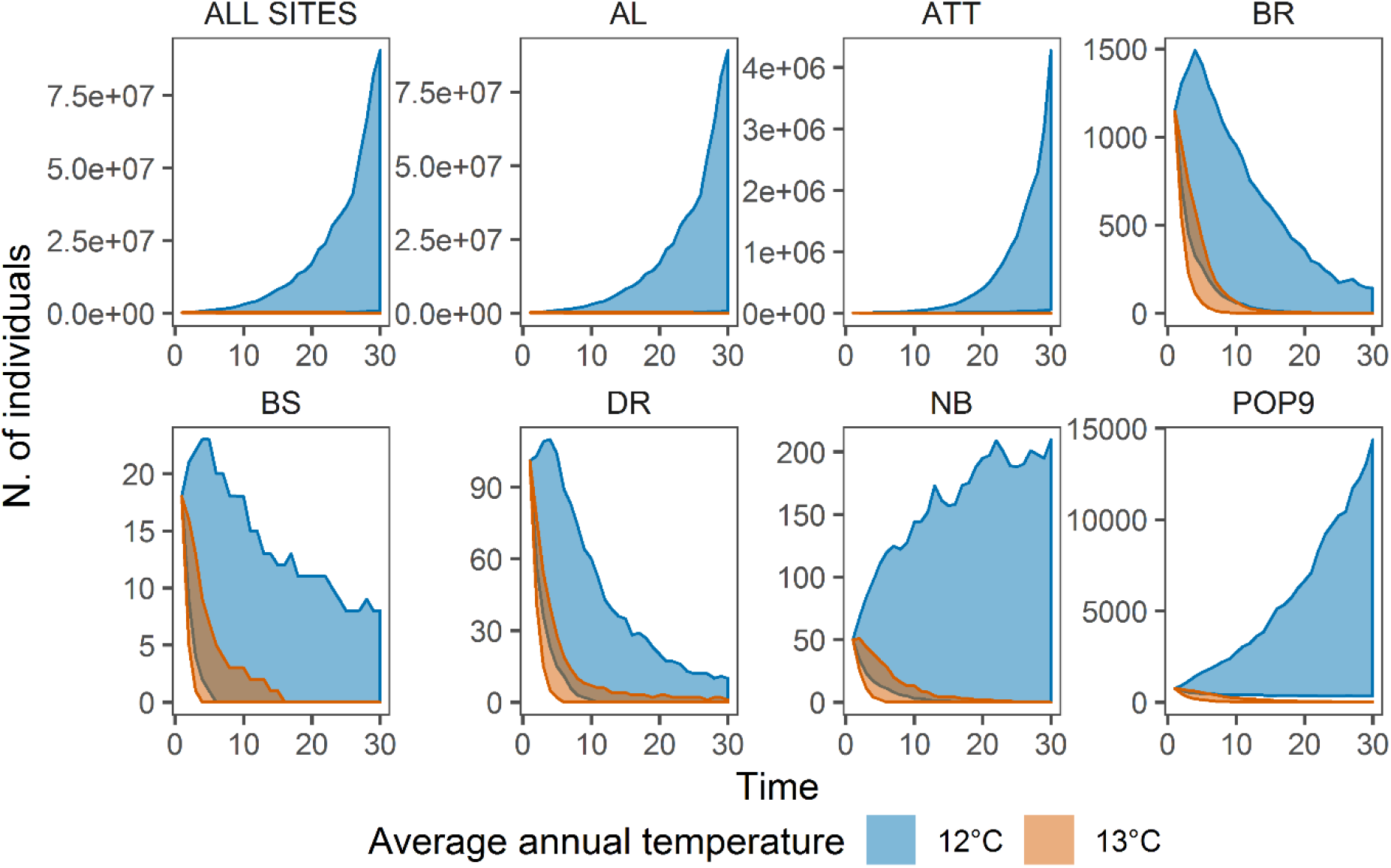
Projected population sizes of *L. tidestromii* assuming historical (12° Celsius) and future average annual temperatures (13° Celsius). Polygons show the lower 2.5^th^ and upper 97.5^th^ percentile of 1000 separate projections. The upper left panel shows the population sizes of all sites at Point Reyes National Seashore; the remaining panels show the population sizes at each one of the seven sites separately. Each simulation was initiated using the population sizes observed in 2018.

## DISCUSSION

Our results demonstrate that climate change is having profound negative effects on the current and future population growth rates of *L. tidestromii*. Within the next 30 years, climate change will result in high probabilities of extinction for four out of the seven populations we analyzed. These high extinction probabilities add to the substantial decrease in total population size. As our analysis considers 7 of the 15 extant populations of the species, including the largest population, it is clear that climate change will substantially deteriorate the conservation status of *L. tidestromii*.

We hypothesize that our results could reflect a general pattern among species evolved in coastal dune habitats experiencing low variation in annual temperatures. Low variation in climatic variables, both temporally and spatially (e.g. across topographic gradients), should decrease the ability of species to respond to climatic changes. This expectation is justified because climatic variation in time and space selects for high genetic variability and high phenotypic plasticity (Nadeau et al. 2017). Dune plants of the pacific coast such as *L. tidestromii* experience low variability in annahope that our study serves as a model ul temperature, and do not occur across large topographic gradients. The standard deviation in annual temperatures at our study populations is only 0.43° Celsius, which is below the 5^th^ percentile of those found across the conterminous United States. Low variation in annual temperatures should be common among Western temperate coasts as a result of Westerly currents. Moreover, dune plants occur only at sea level, and therefore they are not adapted to relatively small topographic gradients. Given these considerations, it might be fruitful to carry out comparative tests on the climatic sensitivity of taxa adapted to dune habitats of mid-latitude Western coasts.

Alternatively, our results for *L. tidestromii* could reflect an isolated case, because climate change can threaten species in any biome as long as their physiological tolerances are exceeded. For example, *L. tidestromii* could be vulnerable because it thrives only within a narrow range of temperatures (Jenouvrier, 2013). One of the reasons for a narrow temperature niche is that the geographic distribution of rare species like *L. tidestromii* could be much smaller than its potential (Svenning and Skov 2004). If *L. tidestromii* did not fill its potential range, its populations could fall, by sheer luck, at sites whose average climate is close to the upper limits of its thermal physiological tolerance. An additional hypothesis for such high climate sensitivity is that the phylogeny of *L. tidestromii* might make this species prone to physiological stress from heat. The productivity of legume crops is very sensitive to heat stress (Liu et al. 2019), a trait which might be shared with wild legumes.

Our results show three advantages of demographic approaches compared to species distribution modeling. First the temporally explicit projections of our demographic models provide highly relevant information to conservation planning and prioritization (Fig. 5). Second, demographic analysis holds mechanistic insights. In our case, we found that the population growth rate of *L. tidestromii* responds to climate mainly through survival. Hence, management actions that increase average individual survival might ameliorate the risks imposed by climate change. Finally, it would be hard or impossible to construct an SDM for *L. tidestromii*. This species occurs in just three locations which differ in average temperature by just 0.12 degrees Celsius. Moreover, the minimum number of sites needed to build an SDM free of statistical artifacts is above 10 (Proosdij et al. 2016). Hence, long-term population or demographic data might be a preferable way to devise climate change forecasts for species that have a geographically restricted range. Some authors suggest that establishing a link between demographic rates and climatic drivers might require 20 to 25 years of data (Teller et al. 2016, Tenhumberg et al. 2018). Instead, our results show that such long data collection might not be needed when a study period spans a large range of climate anomalies (Appendix S5: Fig. S9).

To our knowledge, the evidence we present here is the first example suggesting that climate change should be included as a threat in the status for a species listed under the Endangered Species Act. To date, climate change has been considered in listing actions for four animal species occurring in arctic, alpine, or arid biomes. These animals are the polar bear, the American pika, the American wolverine, and the Gunnison sage-grouse (Blumm and Marienfeld 2013, USFWS 2015). While our results on *L. tidestromii* could be an isolated case, they suggest that the extinction threat posed by climate change might be overlooked in temperate biomes. Thus, we hope that our study serves as a model for other similar forecasting efforts using long-term monitoring data.

**Table 1.** Cross-validation results based on log-likelihood. The models with higher support appear first, those with the least support are last. Model “null” does not contain a climatic predictor. We compare models with four annual climatic anomalies: temperature (tmp), precipitation (ppt), Oceanic Niño Index (oni), and the standardized precipitation evapotranspiration index (spei). Moreover, we compare models with climatic predictors referred to two years: the year of the demographic census (*t*) and that preceding it (*t-1*).

| Survival |  | Growth |  | Flowering |  | Racemes |  |
| --- | --- | --- | --- | --- | --- | --- | --- |
| Model | log_lik_sum | Model | log_lik_sum | Model | log_lik_sum | Model | log_lik_sum |
| tmp<sub>t</sub> | 8320.713 | null | 9643.271 | tmp<sub>t</sub> | 4860.269 | tmp<sub>t</sub> | 20538.58 |
| null | 8339.689 | ppt<sub>t-1</sub> | 9683.304 | null | 5594.391 | null | 20863.18 |
| spei<sub>t-1</sub> | 8378.673 | spei<sub>t</sub> | 9690.057 | oni<sub>t</sub> | 5794.024 | ppt<sub>t</sub> | 20886.95 |
| oni<sub>t</sub> | 8382.177 | ppt<sub>t</sub> | 9696.37 | tmp<sub>t-1</sub> | 5804.951 | spei<sub>to</sub> | 21036.85 |
| tmp<sub>t-1</sub> | 8385.776 | spei<sub>t-1</sub> | 9697.36 | spei<sub>t-1</sub> | 5874.919 | tmp<sub>t-1</sub> | 21236.65 |
| ppt<sub>t</sub> | 8408.783 | oni<sub>t-1</sub> | 9697.919 | ppt<sub>t-1</sub> | 5876.703 | oni<sub>t</sub> | 21281.96 |
| oni<sub>t-1</sub> | 8439.169 | tmp<sub>t-1</sub> | 9700.742 | oni<sub>t-1</sub> | 5882.694 | ppt<sub>t-1</sub> | 21292.39 |
| spei<sub>t</sub> | 8442.731 | oni<sub>t</sub> | 9730.185 | spei<sub>t</sub> | 5888.286 | spei<sub>t-1</sub> | 21326.18 |
| ppt<sub>t-1</sub> | 8494.641 | tmp<sub>t</sub> | 9808.604 | ppt<sub>t</sub> | 5890.526 | oni<sub>t-1</sub> | 21336.45 |

## ACKNOWLEDGMENTS

Funding for this research was provided by the NSF (DEB-0743731), the Alexander von Humboldt foundation, and Washington University in St. Louis. We thank Valentin Ştefan for support with data processing. We thank Dr. Ben Becker at Point Reyes National Seashore for lodging and extensive logistical support, and Lorraine Parsons at Point Reyes National Seashore for long-term collaboration on the monitoring of this species, and sharing site-based knowledge. We thank numerous students for assisting with field data collection, in particular Emily Dangremond and Melissa Patten who worked at least three summers on this project.

## APPENDICES

## Appendix S1: Plant size estimation

We estimated plant size in terms of surface area covered by each individual. However, our way to estimate this are changed before and after 2008. Before 2008 we quantified size counting the number of branches on a plant. After 2008, we quantified the area of each individual approximating it using as an ellipse (e.g. Morris and Doak 2005). The major axis of this ellipse was the longest segment covered by each individual, the minor axis the width perpendicular to the major axis. For data before 2008, we inferred the area of the ellipse by developing a regression that used a sample of plants for which we counted branches and measured major and minor axis length. In total, this regression contained 234 plants (82 measured in 2008, 152 in 2018). We predicted area as a quadratic function of the number of branches (Fig. S1).

**Figure S1.**
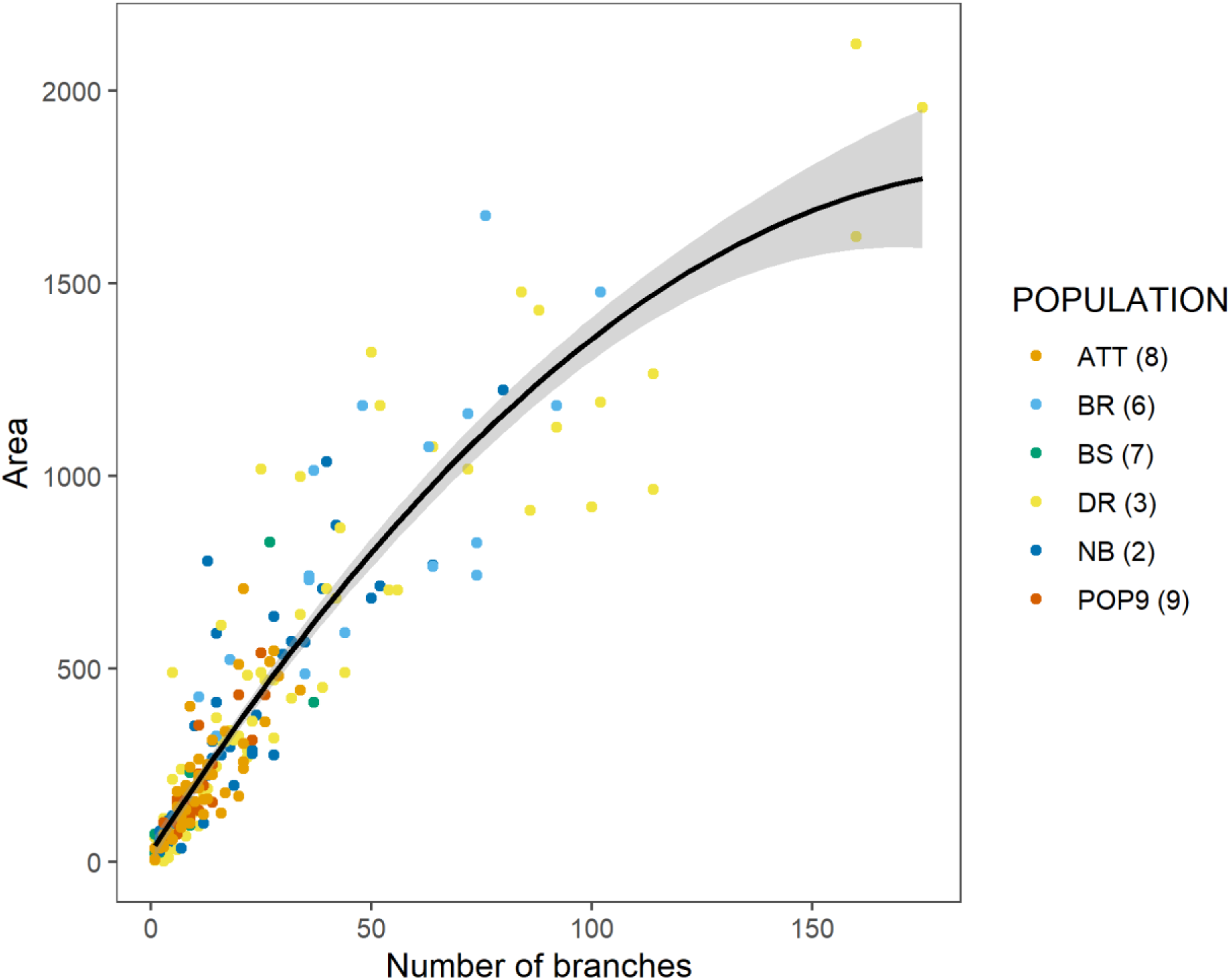
Relationship between the number of branches counted on *L. tidestromii* plants, and their area. The black line represents the mean model prediction of the quadratic linear model that describes the relationship between area and the number of branches. The grey area represents the 95% confidence interval of the mean model prediction. The color of dots refers to data from six separate populations.

## Appendix S2: Vital rates figures

**Figure S2.**
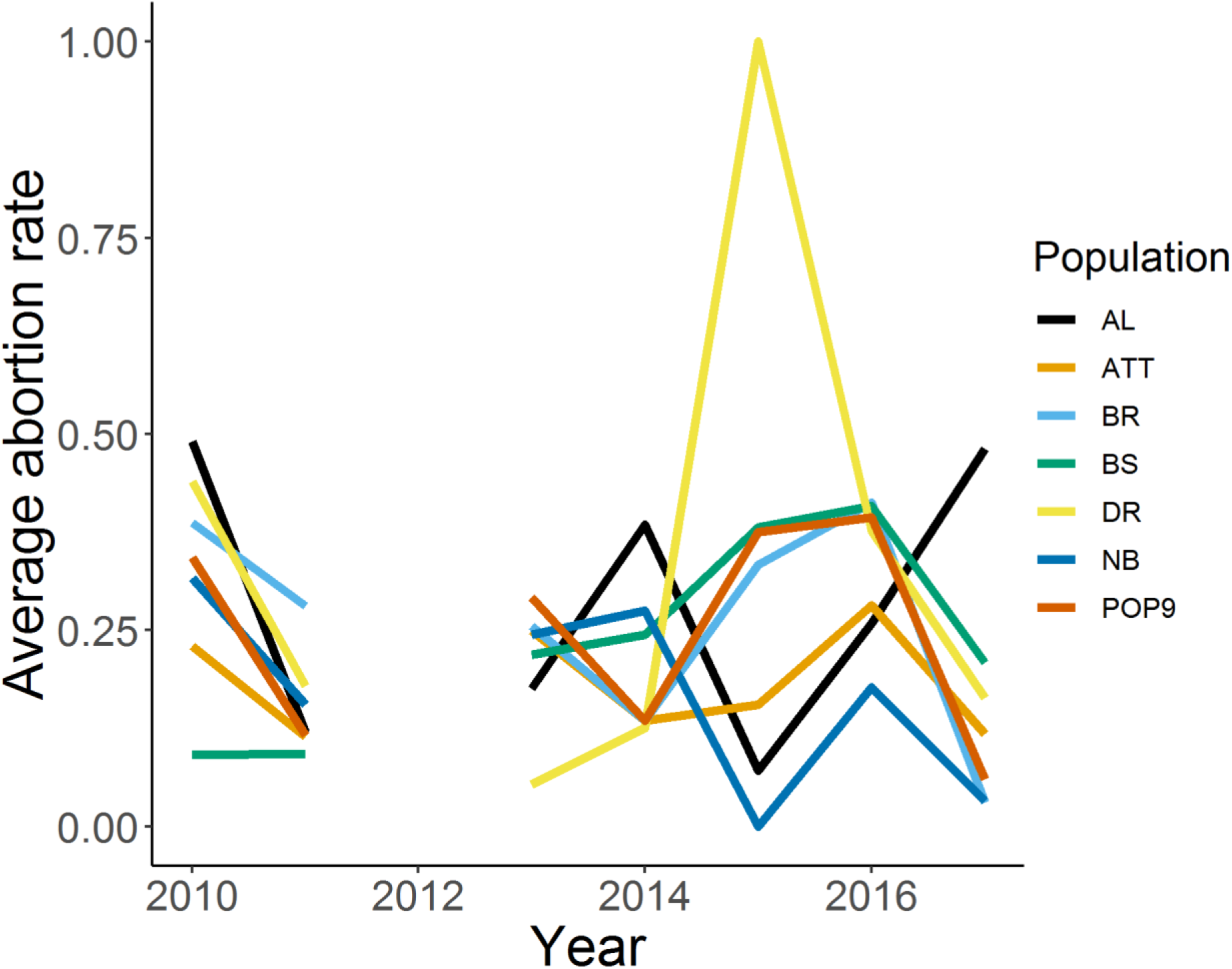
Average abortion rate of inflorescences by year and population.

**Figure S3.**
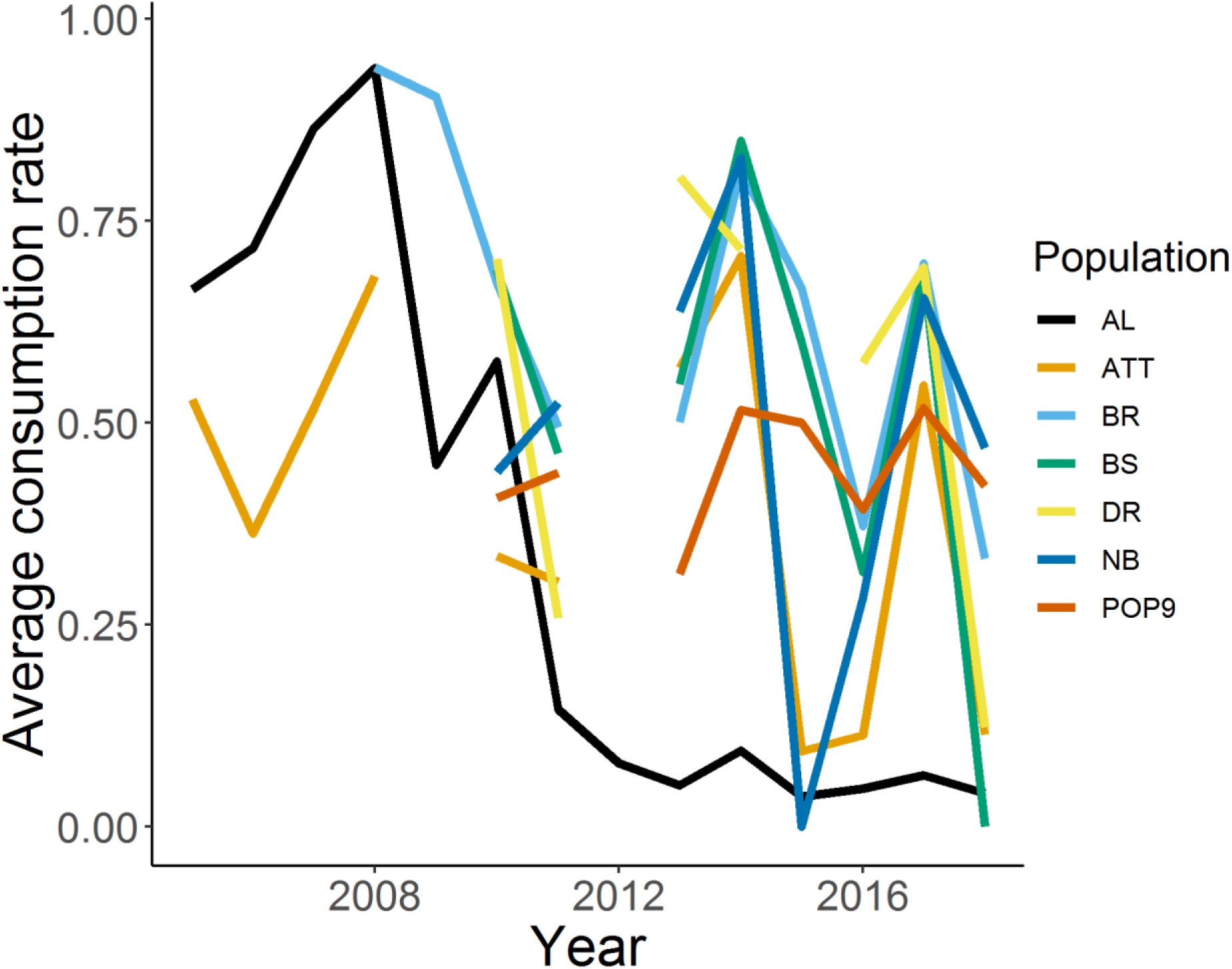
Average consumption rate by year and population.

**Figure S4.**
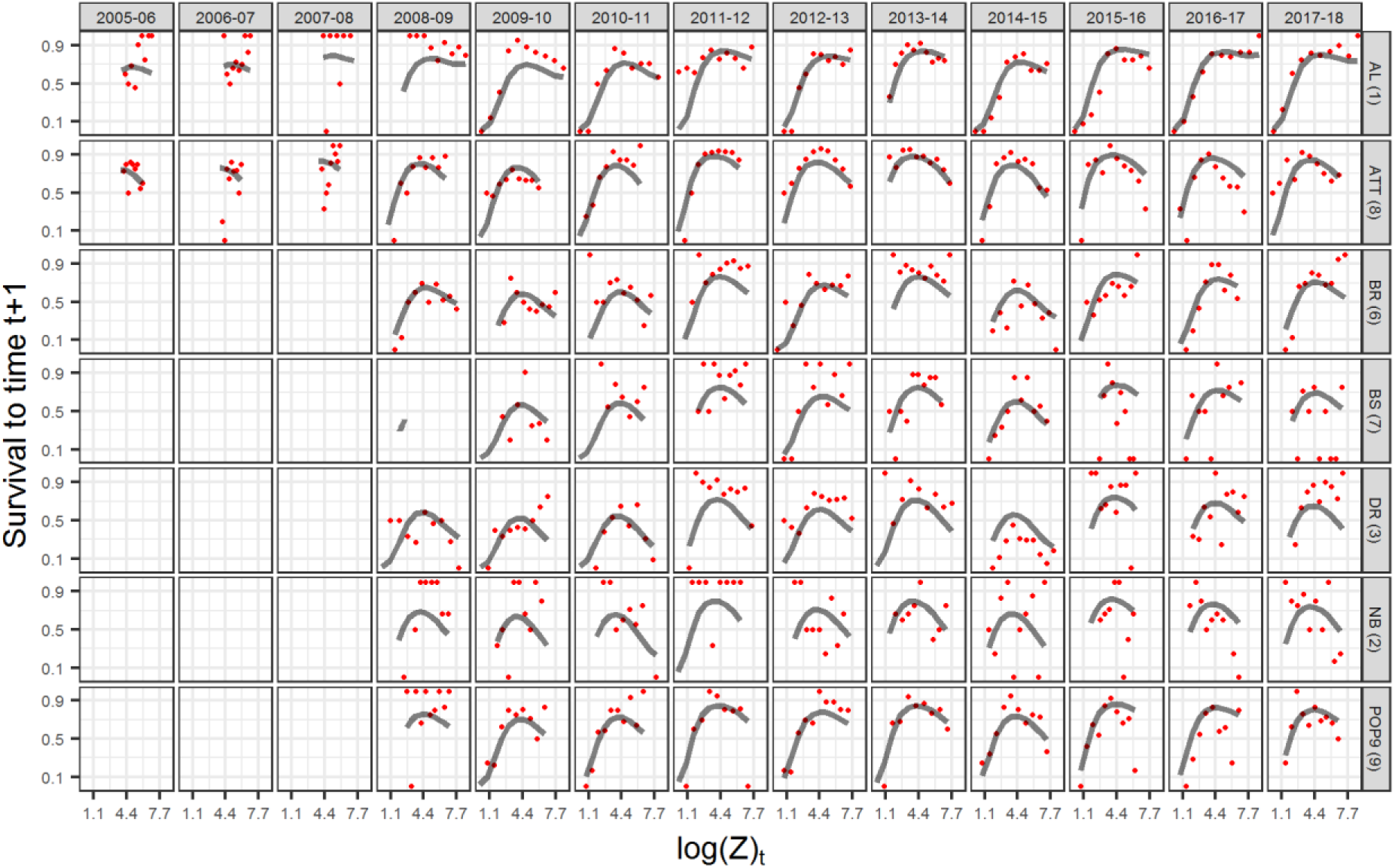
Survival to time t+1 of individuals of size log(Z) at time *t*, by transition year (columns) and population (rows). Red dots show the observed proportion of surviving individuals in ten equally spaced size log(Z) intervals at time *t*. Grey lines show the average size-dependent survival probability predicted by the generalized linear mixed model.

**Figure S5.**
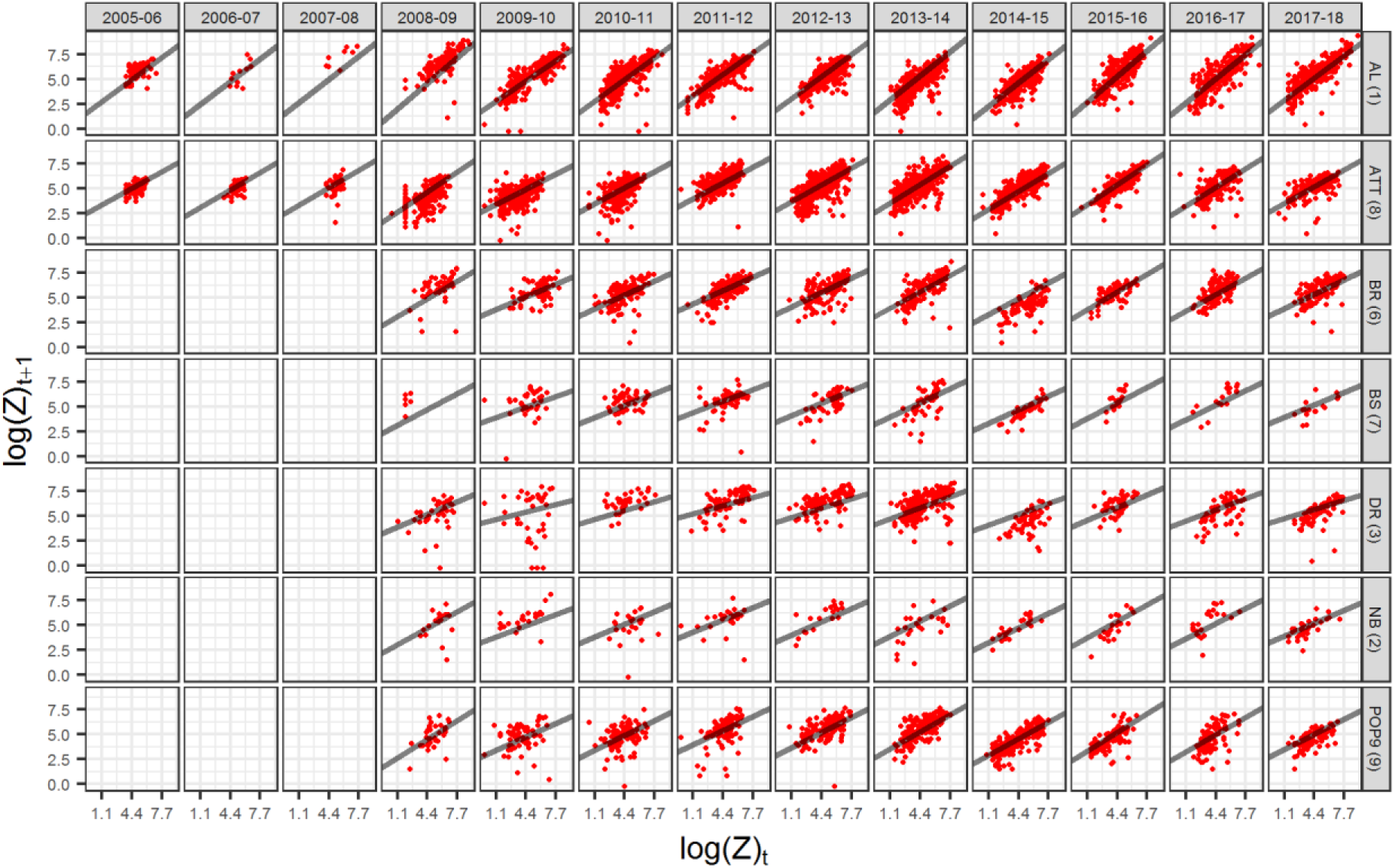
Individual log size (Z) at time t+1 as a function of individual log size (Z) at time t, by transition year (columns) and population (rows). Red dots show individual data points, and grey lines show the prediction of the linear mixed model.

**Figure S6.**
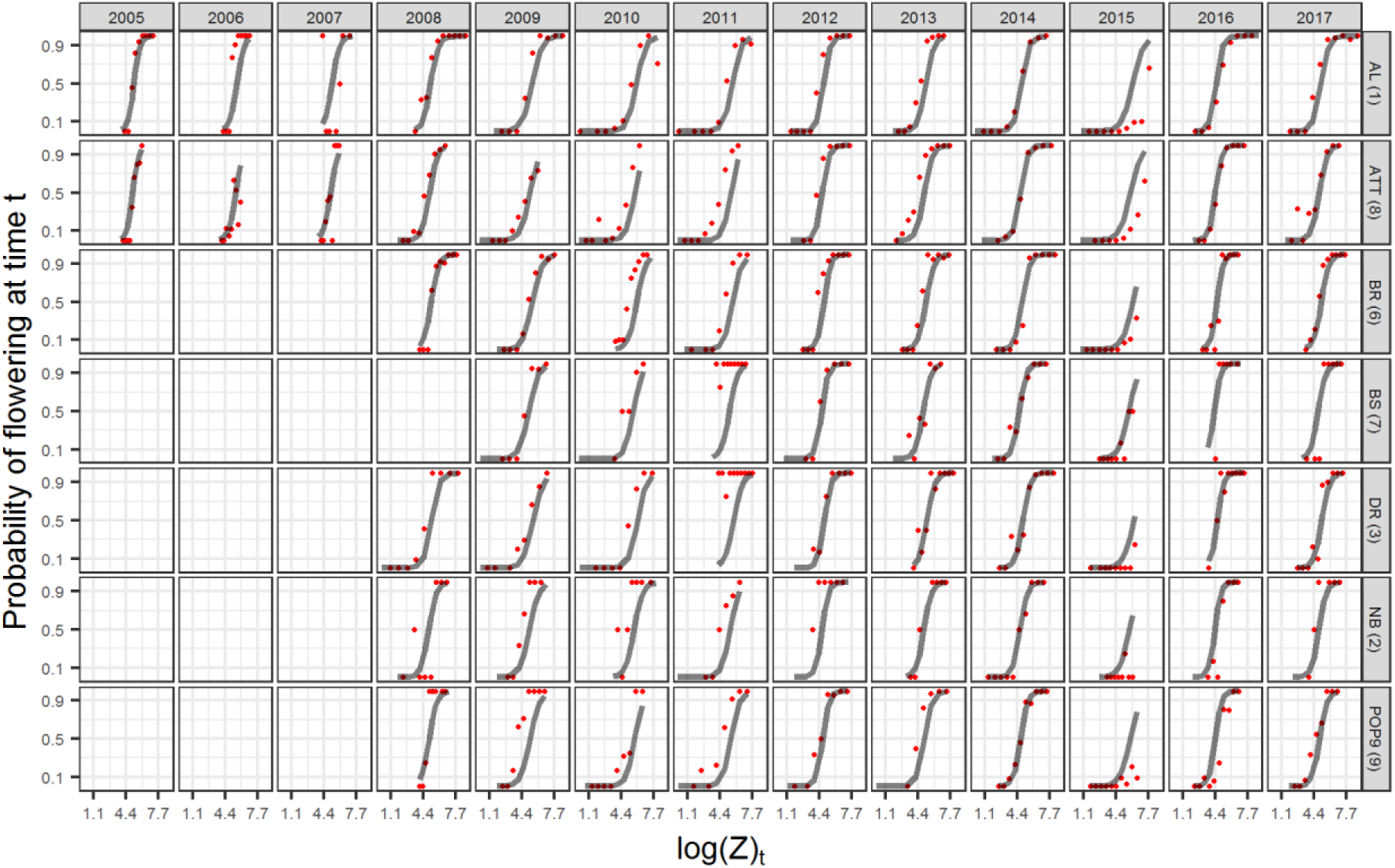
Flowering probability at time *t* based on the log size of individuals (Z) at time *t*, by year (columns) and population (rows). Red dots show the observed proportion of flowering individuals in ten equally spaced intervals of log sizes at time *t*. Grey lines show the average size-dependent flowering probability predicted by the generalized linear mixed model.

**Figure S7.**
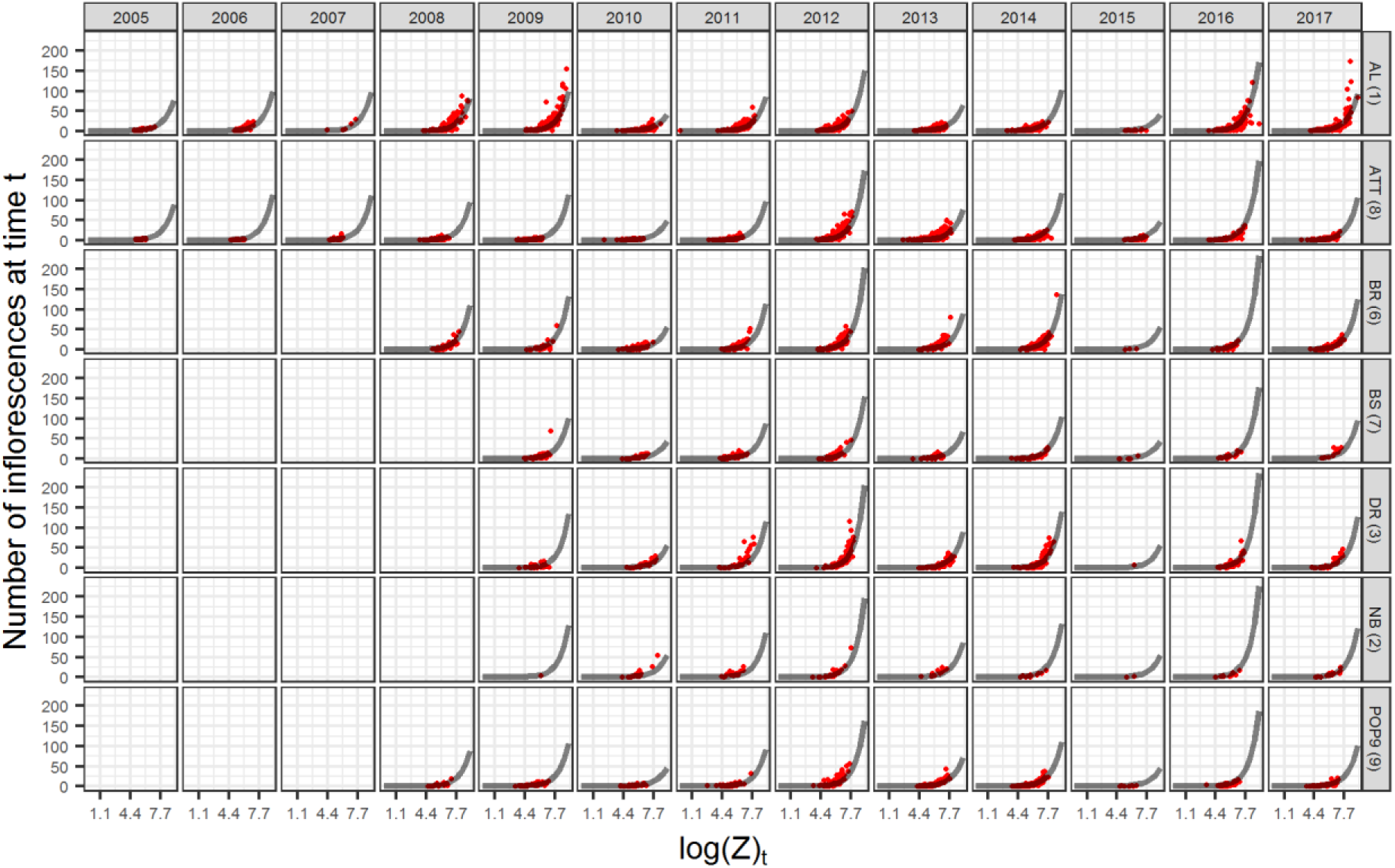
Production of inflorescences by reproductive individuals at time t as a function of individual log size (Z) at time t, by year (columns) and population (rows). Red dots show individual data points, and grey lines show the predictions of the generalized linear mixed model.

**Figure S8.**
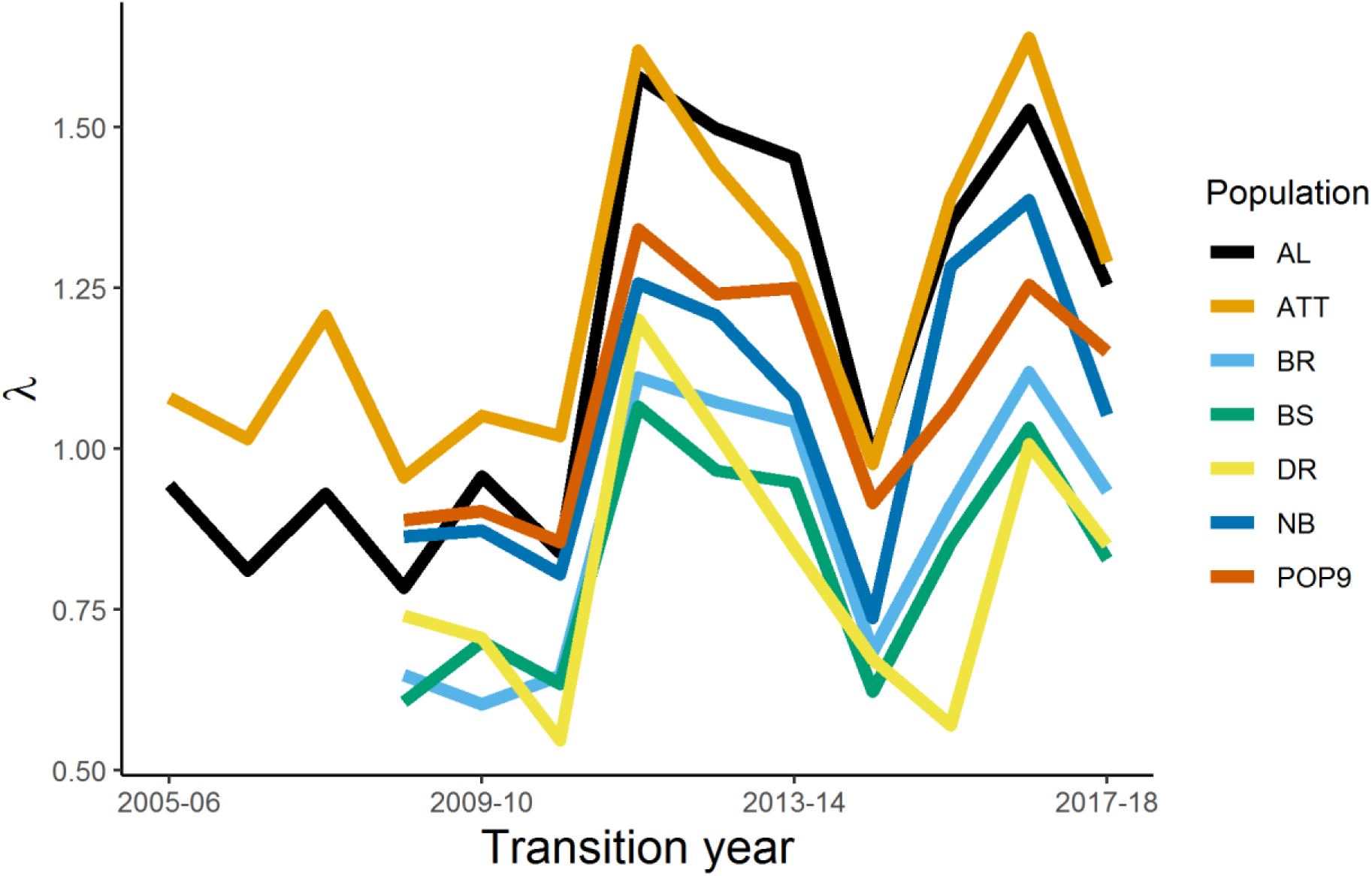
Asymptotic population growth rates (λ) based on transition year and population. λ are calculated using year- and population-specific integral projection models. λ values prior to the 2008-2009 transition are available for two sites only (AL and ATT).

## Appendix S3: Calculation of the germination adjustment factor

We estimated a germination adjustment factor, δ, by combining information on germination trials, on seed production, and on the number of new seedlings. We used data from “known population counts”, which refers to the populations and transition years for which we have counted the total number of individuals and seedlings. First, we estimated the expected number of seeds produced at each population, *p*, and year, *t* (*estimated seedspt*). To estimate the number of seeds, we used year- and population-specific consumption and abortion, and constant values of fruits per inflorescence and seeds per fruit data. We then used the ratio of observed seedlingspt+1 to the estimated seedspt to calculate the recruitment adjustment factor δ:

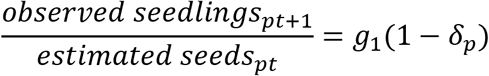

Rearranging,

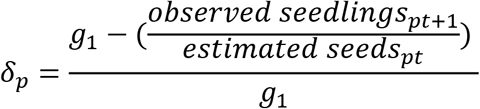

Where *g*_*1*_ is the proportion of seeds germinating in the first year (after one winter) in the germination experiment. In our population models, we apply δ_*p*_ to *g*_2_ and *g*_3_ because calculating δ_*p*_ using *g*_*1*_ provides a conservative estimate of unobserved seed and seedling mortality. Our δ_*p*_ is conservative because the observed seedlings at *t*+1 emerge from dispersed seeds (*g*_*1*_) and from the seed bank (*g*_2_, *g*_3_). We modified germination rates *g*_1_, *g*_2_, and *g*_3_ used in the IPM using δ_*p*_ as:

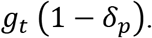

We could only calculate δ_*p*_ for the five populations with known population counts. For population BR, we applied the mean of δ_*p*_ from BS and DR; for AL, we applied the mean of δ_*p*_ from ATT and Pop9. We used these estimates due to the geographic proximity of BR and AL to these smaller populations for which δ could be quantified. We could only calculate δ_*p*_ for the five populations with known population counts. For population BR, we applied the mean of δ_*p*_ from BS and DR; for AL, we applied the mean of δ_*p*_ from ATT and Pop9. We used these estimates due to the geographic proximity of BR and AL to these smaller populations for which δ could be quantified.

## Appendix S4: Individual based model

Our individual based model simulated each probabilistic event in the life cycle of *L. tidestromii* (Appendix S1: Fig. S3) using an appropriate statistical distribution. Our probabilistic events were linked to survival, growth, probability of flowering, production of inflorescences, the abortion of inflorescences, the consumption of inflorescences, and the germination of seeds. Starting from the production of flowers, each individual flowered based on a Bernoulli process depending on its individual size, so that:

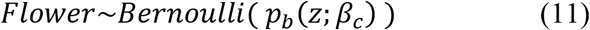

Where *Flower* is either a 0, if an individual failed to flower, or 1, if an individual flowered.

*p*_*b*_(*z*;*β*_*c*_) shows that the probability of reproduction depends on individual size, *z*, and on the climate anomaly. If an individual reproduces, its production of inflorescences depends on a negative binomial distribution, so that

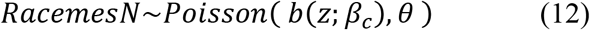

Which is also a size- and climate-dependent process. The number of inflorescences would decrease because of abortion following a binomial process

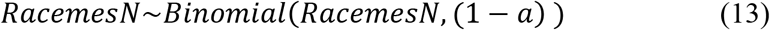

where 1-*a* is the probability of an inflorescence not aborting. The remaining inflorescences could still be consumed by predators,

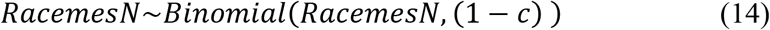

where 1-*c* is the probability of not being consumed (the probability of experiencing pre-dispersal predation). The seeds produced by the surviving inflorescences on each individual could then germinate based on a binomial process, so that:

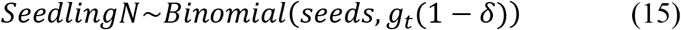

Where SeedlingN is the number of seedlings emerging in each population, and *gt* is one of the germination rates associated with seeds that germinate in the year of the transition (g_0_), the year after (g_1_), and two years after (g_2_). Finally, individuals could survive based on a size- and climate-dependent Bernoulli process

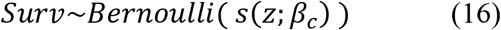

and, in case they survived, they could grow based on a normal process:

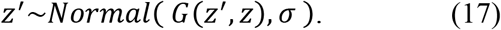

## Appendix S5: Climatic anomalies figure

**Figure S9.**
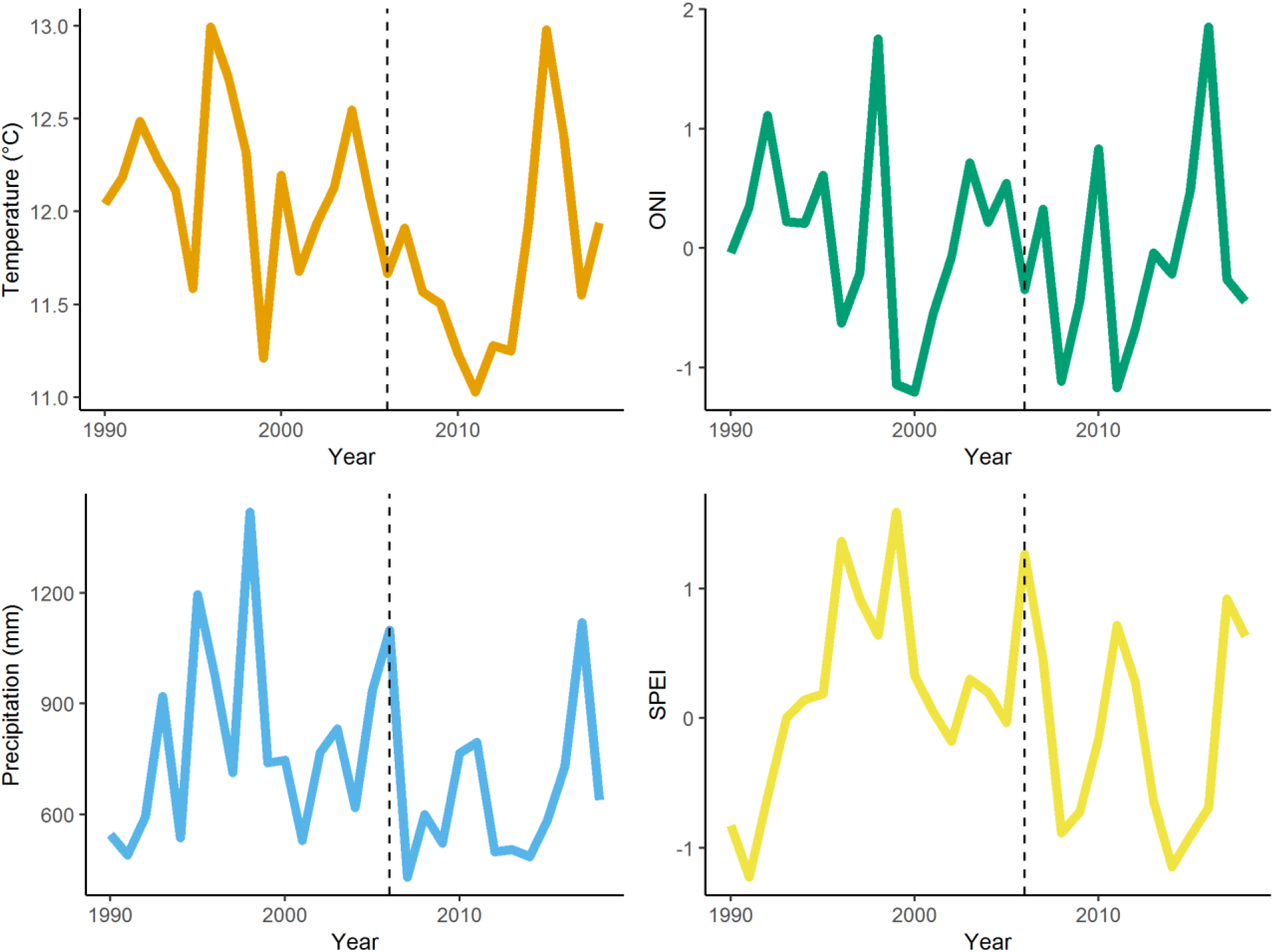
Annual climate values of temperature, precipitation, Oceanic Niño Index (ONI), and standardized aridity index (SPEI) between 1990 and 2018. The dashed lines denote the beginning of *L. tidestromii* demographic censuses.

## Appendix S6: Elasticity analysis

**Figure S10.**
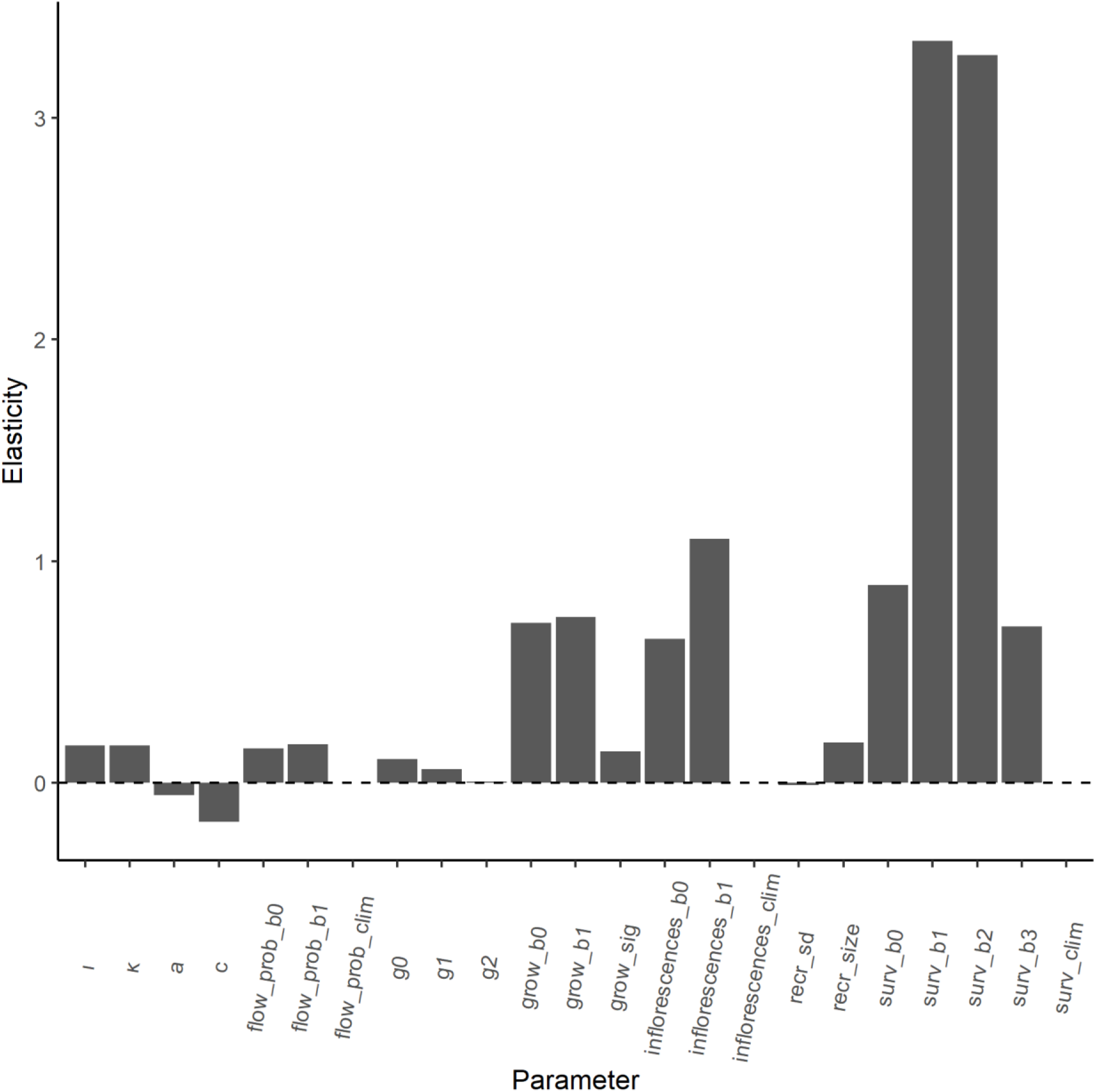
Elasticity of asymptotic population growth rate (λ) to the parameters of the IPM. Parameters refer to the average values across years and populations. The elasticity is a proportional sensitivity, meaning that it quantifies by what percentage λ will change after a percent change in each parameter. Thus, elasticity facilitates the comparison among parameters. The parameter names for size-dependent processes show the effect of annual temperature anomaly (_clim), the intercept of linear models (_b0), and the linear (_b1), quadratic (_b2), and cubic (_b3) effects of size. Parameters “recr_size” and “recr_sd” refers to the mean size and standard deviation of recruits. The remaining parameter names are identical to those in equations 7–9. Note that, unlike the elasticity of the IPM kernel elements, the elasticity of lower level parameters can exceed one.

## REFERENCES

Baldwin, B. G., D. H. Goldman, D. J. Keil, R. Patterson, T. J. Rosatti, and D. H. Wilken. 2012. The Jepson Manual: Vascular Plants of California. University of California Press.

Barbier, E. B. 2015. Valuing the storm protection service of estuarine and coastal ecosystems. Ecosystem Services 11:32–38.

Bates, D., M. Mächler, B. Bolker, and S. Walker. 2015. Fitting Linear Mixed-Effects Models Using {lme4}. Journal of Statistical Software 67.

Beguería, S., and S. Vicente-Serrano. 2017. SPEI: Calculation of the Standardised Precipitation-Evapotranspiration Index.

Blumm, M. C., and K. B. Marienfeld. 2013. Endangered Species Act Listings and Climate Change: Avoiding the Elephant in the Room Symposium. Animal Law:277–310.

Burnham, K. P., and D. R. Anderson. 2002. Model Selection and Multimodel Inference: A Practical Information-Theoretic Approach. Second edition. Springer-Verlag, New York.

Campbell, D. R. 2019. Early snowmelt projected to cause population decline in a subalpine plant. Proceedings of the National Academy of Sciences 116:12901–12906.

Caswell, H. 2000. Prospective and Retrospective Perturbation Analyses: Their Roles in Conservation Biology. Ecology 81:619–627.

Caswell, H. 2001. Matrix population models. Massachusetts: Sinauer Associates.

Cogoni, D., E. Sulis, G. Bacchetta, and G. Fenu. 2019. The unpredictable fate of the single population of a threatened narrow endemic Mediterranean plant. Biodiversity and Conservation 28:1799–1813.

CPC. (n.d.). https://origin.cpc.ncep.noaa.gov/products/analysis_monitoring/ensostuff/ONI_v5.php.

Daly, C., R. P. Neilson, and D. L. Phillips. 1994. A Statistical-Topographic Model for Mapping Climatological Precipitation over Mountainous Terrain. Journal of Applied Meteorology 33: 140–158.

Dangremond, E. M., E. A. Pardini, and T. M. Knight. 2010a. Apparent competition with an invasive plant hastens the extinction of an endangered lupine. Ecology 91:2261–2271.

Dangremond, E. M., E. A. Pardini, and T. M. Knight. 2010b. Apparent competition with an invasive plant hastens the extinction of an endangered lupine 91:11.

Díaz, S., J. Settele, E. Brondízio, H. T. Ngo, M. Guèze, J. Agard, A. Arneth, P. Balvanera, K. Brauman, R. T. Watson, I. A. Baste, A. Larigauderie, P. Leadley, U. Pascual, B. Baptiste, S. Demissew, L. Dziba, G. Erpul, A. Fazel, M. Fischer, A. María, M. Karki, V. Mathur, T. Pataridze, I. S. Pinto, M. Stenseke, K. Török, and B. Vilá. 2019. Summary for policymakers of the global assessment report on biodiversity and ecosystem services of the Intergovernmental Science-Policy Platform on Biodiversity and Ecosystem Services:45.

Doak, D. F., and W. F. Morris. 2010. Demographic compensation and tipping points in climate-induced range shifts. Nature 467:959–962.

Ehrlén, J., and W. F. Morris. 2015. Predicting changes in the distribution and abundance of species under environmental change. Ecology Letters 18:303–314.

Elith, J., and J. R. Leathwick. 2009. Species Distribution Models: Ecological Explanation and Prediction Across Space and Time. Annual Review of Ecology, Evolution, and Systematics 40:677–697.

Ellner, S. P., D. Z. Childs, and M. Rees. 2016. Data-driven Modelling of Structured Populations: A Practical Guide to the Integral Projection Model. Springer International Publishing.

Evens, J. G. 2008. natural history of the Point Reyes Peninsula. University of California Press.

Feagin, R. A., D. J. Sherman, and W. E. Grant. 2005. Coastal erosion, global sea-level rise, and the loss of sand dune plant habitats. Frontiers in Ecology and the Environment 3:359–364.

Glenn, E. M., R. G. Anthony, and E. D. Forsman. 2010. Population trends in northern spotted owls: Associations with climate in the Pacific Northwest. Biological Conservation 143:2543–2552.

Gneiting, T., and A. E. Raftery. 2007. Strictly Proper Scoring Rules, Prediction, and Estimation. Journal of the American Statistical Association 102:359–378.

Hunter, C. M., H. Caswell, M. C. Runge, E. V. Regehr, S. C. Amstrup, and I. Stirling. 2010. Climate change threatens polar bear populations: a stochastic demographic analysis. Ecology 91:2883–2897.

Iler, A. M., A. Compagnoni, D. W. Inouye, J. L. Williams, P. J. CaraDonna, A. Anderson, and T. E. X. Miller. 2019. Reproductive losses due to climate change-induced earlier flowering are not the primary threat to plant population viability in a perennial herb. Journal of Ecology 107:1931–1943.

IPCC. 2014. Summary for policymakers. In: Climate Change 2014: Impacts, Adaptation, and Vulnerability. Part A: Global and Sectoral Aspects. Contribution of Working Group II to the Fifth Assessment Report of the Intergovernmental Panel on Climate Change. Cambridge University Press, Cambridge, UK and New York, NY, USA.

Jenouvrier, S. 2013. Impacts of climate change on avian populations. Global Change Biology 19:2036–2057.

Jenouvrier, S., H. Caswell, C. Barbraud, M. Holland, J. Strœve, and H. Weimerskirch. 2009. Demographic models and IPCC climate projections predict the decline of an emperor penguin population. Proceedings of the National Academy of Sciences 106:1844–1847.

Jonzén, N., T. Pople, J. Knape, and M. Sköld. 2010. Stochastic demography and population dynamics in the red kangaroo Macropus rufus. Journal of Animal Ecology 79:109–116.

Keenan, T., J. M. Serra, F. Lloret, M. Ninyerola, and S. Sabate. 2011. Predicting the future of forests in the Mediterranean under climate change, with niche- and process-based models: CO2 matters! Global Change Biology 17:565–579.

Lande, R. 1993. Risks of Population Extinction from Demographic and Environmental Stochasticity and Random Catastrophes. The American Naturalist 142:911–927.

Liu, Y., J. Li, Y. Zhu, A. Jones, R. J. Rose, and Y. Song. 2019. Heat Stress in Legume Seed Setting: Effects, Causes, and Future Prospects. Frontiers in Plant Science 10.

Luger, M. I. 1991. The economic value of the coastal zone:23.

Maron, J. L. 1998. Insect Herbivory Above- and Belowground: Individual and Joint Effects on Plant Fitness. Ecology 79:1281–1293.

Martínez, M. L., A. Intralawan, G. Vázquez, O. Pérez-Maqueo, P. Sutton, and R. Landgrave. 2007. The coasts of our world: Ecological, economic and social importance. Ecological Economics 63:254–272.

Morris, W. F., and D. F. Doak. 2005. How General Are the Determinants of the Stochastic Population Growth Rate Across Nearby Sites? Ecological Monographs 75:119–137.

Paniw, M., N. Maag, G. Cozzi, T. Clutton-Brock, and A. Ozgul. 2019. Life history responses of meerkats to seasonal changes in extreme environments. Science 363:631–635.

Pardini, E. A., K. E. Vickstrom, and T. M. Knight. 2015. Early Successional Microhabitats Allow the Persistence of Endangered Plants in Coastal Sand Dunes. PLOS ONE 10:e0119567.

Pereira, H. M., P. W. Leadley, V. Proença, R. Alkemade, J. P. W. Scharlemann, J. F. Fernandez-Manjarrés, M. B. Araújo, P. Balvanera, R. Biggs, W. W. L. Cheung, L. Chini, H. D. Cooper, E. L. Gilman, S. Guénette, G. C. Hurtt, H. P. Huntington, G. M. Mace, T. Oberdorff, C. Revenga, P. Rodrigues, R. J. Scholes, U. R. Sumaila, and M. Walpole. 2010. Scenarios for Global Biodiversity in the 21st Century. Science 330:1496–1501.

Pisanu, S., E. Farris, R. Filigheddu, and M. B. García. 2012. Demographic effects of large, introduced herbivores on a long-lived endemic plant. Plant Ecology 213:1543–1553.

Proosdij, A. S. J. van, M. S. M. Sosef, J. J. Wieringa, and N. Raes. 2016. Minimum required number of specimen records to develop accurate species distribution models. Ecography 39:542–552.

Sæther, B.-E., J. Tufto, S. Engen, K. Jerstad, O. W. Røstad, and J. E. Skåtan. 2000. Population Dynamical Consequences of Climate Change for a Small Temperate Songbird. Science 287:854–856.

Schurr, F. M., J. Pagel, J. S. Cabral, J. Groeneveld, O. Bykova, R. B. O’Hara, F. Hartig, W. D. Kissling, H. P. Linder, G. F. Midgley, B. Schröder, A. Singer, and N. E. Zimmermann. 2012. How to understand species’ niches and range dynamics: a demographic research agenda for biogeography. Journal of Biogeography 39:2146–2162.

Svenning, J.-C., and F. Skov. 2004. Limited filling of the potential range in European tree species. Ecology Letters 7:565–573.

Teller, B. J., P. B. Adler, C. B. Edwards, G. Hooker, and S. P. Ellner. 2016. Linking demography with drivers: climate and competition. Methods in Ecology and Evolution 7:171–183.

Tenhumberg, B., E. E. Crone, S. Ramula, and A. J. Tyre. 2018. Time-lagged effects of weather on plant demography: drought and Astragalus scaphoides. Ecology 99:915–925.

Tews, J., and F. Jeltsch. 2004. Modelling the impact of climate change on woody plant population dynamics in South African savanna. BMC Ecology 4:17.

USFWS. 1998. Seven coastal plants and the Myrtle’s silverspot butterfly recovery plan. USFWS, Portland, Oregon, USA.

USFWS. 2009. Lupinus tidestromii (Clover lupine) 5-Year Review: Summary and Evaluation. Sacramento Fish and Wildlife Office, Sacramento, California, USA.

USFWS. 2015. Polar Bear (Ursus maritimus) Conservation Management Plan, Draft. U.S. Fish and Wildlife, Region 7, Anchorage, Alaska.

USGCRP. 2018. Fourth National Climate Assessment. U.S. Global Change Research Program, Washington, DC, USA.

Vicente-Serrano, S. M., S. Beguería, and J. I. López-Moreno. 2009. A Multiscalar Drought Index Sensitive to Global Warming: The Standardized Precipitation Evapotranspiration Index. Journal of Climate 23:1696–1718.

Vose, R. S., D. R. Easterling, K. E. Kunkel, A. N. LeGrande, and M. F. Wehner. 2017. Temperature Changes in the United States. Climate Science Special Report: Fourth National Climate Assessment, Volume I. U.S. Global Change Research Program.

Wang, T., A. Hamann, D. Spittlehouse, and C. Carroll. 2016. Locally Downscaled and Spatially Customizable Climate Data for Historical and Future Periods for North America. PLOS ONE 11:e0156720.

Wang, T., A. Hamann, D. Spittlehouse, and C. Carroll. (n.d.). ClimateNA_MAP -- An Interactive Plantform for Visualization and Data Access.

## REFERENCES

Morris, W.F. & Doak, D.F. (2005). How General Are the Determinants of the Stochastic Population Growth Rate Across Nearby Sites? Ecological Monographs, 75, 119–137.

